# Auxin promotion of seedling growth via ARF5 is dependent on the brassinosteroid-regulated transcription factors BES1 and BEH4

**DOI:** 10.1101/664144

**Authors:** Anahit Galstyan, Jennifer L Nemhauser

**Affiliations:** Department of Biology, University of Washington, Seattle, WA 98195-1800, USA

**Author notes:** Max Planck Institute for Plant Breeding Research, Carl-von-Linné-Weg 10, 50829, Cologne, Germany (Anahit Galstyan).

**Keywords:** auxin, brassinosteroids, promoter architecture, growth related genes, seedling development, *Arabidopsis thaliana*, transcriptional modules

## Abstract

Seedlings must continually calibrate their growth in response to the environment. Auxin and brassinosteroids (BRs) are plant hormones that work together to control growth responses during photomorphogenesis. We used our previous analysis of promoter architecture in an auxin and BR target gene to guide our investigation into the broader molecular bases and biological relevance of transcriptional co-regulation by these hormones. We found that the auxin-regulated transcription factor AUXIN RESPONSIVE FACTOR 5 (ARF5) and the brassinosteroid-regulated transcription factor BRI1-EMS-SUPPRESOR 1/BRASSINOZOLERESISTANT 2 (BES1) co-regulated a subset of growth promoting genes via conserved bipartite *cis*-regulatory elements. Moreover, ARF5 binding to DNA could be enriched by increasing BES1 levels. The evolutionary loss of bipartite elements in promoters results in loss of hormone responsiveness. We also identified another member of the BES1/BZR1 family called BEH4 that acts partially redundantly with BES1 to regulate seedling growth. Double mutant analysis showed that BEH4 and not BZR1 were required alongside BES1 for normal auxin response during early seedling development. We propose that an ARF5-BES1/BEH4 transcriptional module acts to promote growth via modulation of a diverse set of growth-associated genes.

## Introduction

Plants must adapt their form to survive a complex and changing environment. The extensive molecular interplay between external (i.e., light and temperature) and internal (i.e., circadian clock) signals allows for a high degree of developmental plasticity. Light-directed seedling growth (photomorphogenesis) is one of the best-characterized examples of a highly dense regulatory network. Small molecule hormones are critical for relaying information about the light environment, as well as a diverse set of additional metabolic, environmental and developmental cues [1, 2]. Hormones like auxin and brassinosteroids (BRs) play a central role in coordinating growth during photomorphogenesis.

Plants with defective responses to auxin or BRs show an array of phenotypes of light-grown plants even when grown in the dark [3]. The signaling pathways downstream of auxin and BRs are distinct. Auxin binds to the TRANSPORT INHIBITOR RESPONSE 1/AUXIN SIGNALING F-BOX 1–5 (TIR1/AFB) family of F-box receptors and triggers the ubiquitination and degradation of AUXIN/INDOLE-3-ACETIC ACID (Aux/IAA) co-repressors. Loss of the Aux/IAAs activates AUXIN RESPONSE FACTORs (ARFs) to regulate gene expression [4–6]. ARFs bind to an auxin response element (AuxRE, TGTCTC) and related *cis*-elements in target promoters [7–11]. In contrast, BRs bind and activate the BRASSINOSTEROID-INSENSITIVE1 (BRI1)-associated receptor complex at the plasma membrane. A phospho-relay cascade culminates in dephosphorylated and nuclear-localized transcription factors, including BRI1-EMS-SUPPRESOR1/ BRASSINOZOLERESISTANT2 (BES1/BZR2, hereafter BES1) and BZR1 [12–14]. BES1 and BZR1 regulate gene expression by binding to both E-box (CANNTG) and BRRE (CGTG(T/C)G) *cis*-elements in target promoters [15, 16].

Genetic, physiological, and genomic analyses demonstrate molecular and physiological responses of auxin and BRs are interdependent. BRs promote auxin transport, hence altering overall auxin distribution within the plant [17, 18]. BRs also regulate expression of genes involved in the core auxin response [19–21] and a BR-regulated kinase targets members of the ARF family [22–25]. Auxin stimulates *de novo* BR biosynthesis by directly regulating expression of *DWARF4* (DWF4), a BR biosynthetic enzyme [26, 27]. BRI1 is a direct target of activator ARF5/MONOPTEROS (hereafter, ARF5) [28]. In addition ARF6 and ARF7 were shown to interact with BZR1 to regulate shared target genes [29]. Previously, we demonstrated that a bipartite element in the promoter of *SAUR15* gene that includes a type of E-box called a HUD element (CACATG) and a variant of the AuxRE (TGTCT) are bound by ARF5 and BES1, and that binding by both transcription factors is required for induction of expression by either hormone [30]. In this work, we expanded this study to include other growth-associated genes with predicted bipartite elements in their promoters. We found that BES1 sensitizes hypocotyl response to auxin by enhancing ARF5 binding to shared target promoters. The evolutionary loss of the conserved promoter architecture with bipartite elements results in loss of hormone responsiveness. BEH4, a previously uncharacterized paralog of BES1, was found to act redundantly with BES1 as a major regulator of seedling growth. We propose a model where shared promoter architecture facilitates a coordinated and highly responsive growth controlling module encompassing genes from diverse families.

## Materials and Methods

### Plant materials and growth conditions

The wild type is Arabidopsis thaliana ecotype Col-0 except *beh1-2* and *beh2-1* that are in Col-3 background. *bes1-D* [12], *bzr1-D* [58], *bin2-D* [59], *lng1-3* [46], *xth17* [52], *pif7-2* [60], ARF5_PRO_::ARF5:GFP [31], BES1_PRO_::BES1:GFP [12], XTH19_PRO-1.1kb_::GUS and XTH19_PRO-0.3kb_::GUS [34] were previously described. Single T-DNA insertion lines: *bes1-2* (WiscDsLox 246D02), *bzr1-2* (GABI-Kat 857E04), *beh1-2* (SAIL_40_D04, Col-3 background), *beh2-1* (SAIL_76_B06, Col-3 background), *beh3-1* (SALK_017577), *beh4-1* (SAIL_750_F08) and double mutants: *bes1-2beh4-1* and *bes1-2bzr1-2* are described lines [33]. For detailed information on genotyping methods, primers and generation of double mutants see Supplementary data.

For seed production and crosses, plants were grown in a growth chamber under LD conditions. Seeds were surface sterilized (20 min in 70% ethanol, 0.01% Triton X-100, followed by a rinse in 95% ethanol) for all the physiological and molecular analyses. For hypocotyl and GUS assays, sterilized seeds were suspended in water and sown individually on plates containing 0.5x Linsmaier and Skoog (LS) (LSP03, Caisson Laboratories, Inc., http://www.caissonlabs.com/) with 0.8% phytoagar (40100072-1, Plant Media: bioWorld, http://www.plantmedia.com/), and stratified in the dark at 4°C for 3 days. Plates were placed vertically in a Percival E-30B growth chamber set at 20°C in 30 µmol m^−2^ s^−1^ of photosynthetically active radiation white light with short-day conditions (8 h light, 16 h dark). For gene expression and ChIP assays, sterilized seeds were suspended in 0.1% agar (BP1423, Fisher Scientific, http://www.fisher.co.uk/), spotted on plates containing 0.5x LS with 0.8% phytoagar, stratified in the dark at 4°C for 3 days and grown horizontally as described above.

### Chemical treatments

To prepare stock solutions brassinosteroid (brassinolide, 101, Chemiclones, Inc., www.chemiclones.com), IAA (705490, PlantMedia.com) and picloram (P5575, Sigma) were dissolved in dimethyl sulfoxide and diluted directly into plate medium to 1µM/0,5µM, 50µM and 5µM concentration respectively. Stock solutions were kept at –20°C until use.

### Accession numbers

Sequence data from this article can be found in the Arabidopsis Genome Initiative or GenBank/EMBL databases under the following accession numbers: *BES1* (At1g19350), *BZR1* (At1g75080), *BEH1* (At3g50750), *BEH2* (At4g36780), *BEH3* (At4g18890), *BEH4* (At1g78700), *ARF5* (At1g19850), *IAA6* (At1g52830), *XTH14* (At4g25820), *XTH17* (At1g65310), *XTH18* (At4g30280), *XTH19* (At4g30290), *XTH26* (At4g28850), *XTH31* (At3g44990), *XTH32* (At2g36870), *LNG1* (At5g15580), *PIF7* (At5g61270), *ACT2* (At3g18780)and housekeeping gene-HK (At1g13320).

### Hypocotyl measurements

Seedlings were grown vertically on square plates with 0.5xLS media supplemented with 80% ethanol (mock), 5µM picloram or 0.5µM BL under abovementioned conditions. Plates were scanned using EPSON Perfection V5000 scanner every 24h 2 days after germination. Generated images were used to measure hypocotyl length. The National Institutes of Health ImageJ software (rsb.info.nih.gov) was used on digital images to measure the length of different organs of the seedlings, as indicated elsewhere [61]. At least 15 seedlings were used for each data point, experiments were repeated 3-5 times and a representative one is shown. Statistical analyses of the data (T-test and two-way ANOVA) were performed using GraphPad Prism version 4.00 for Windows (www.graphpad.com).

### Gene expression analysis by RT-qPCR

Seedlings were grown vertically on 0.5xLS plates under abovementioned conditions. Expression analyses were performed on seedlings collected at dawn on day 7 (D8) grown in SD. Only the expression analysis of BES1-BZR1-BEH family members in single mutants was performed on dark grown 7-day-old seedlings (D8). Hormone treatments were done 3h prior to collection in liquid 0.5x LS media supplemented with equal volumes of 80% ethanol (mock), IAA (to final concentration 50µM) or BL (to final concentration 1µM). All samples were immediately frozen in liquid nitrogen and stored at −80°C until processing. Total RNA was extracted from 100 mg of whole seedling tissue using the Spectrum Plant Total RNA Kit (Sigma, STRN50), treated with DNaseI on columns (Qiagen, DNASE70) and lµg of eluted RNA was used for complementary DNA (cDNA) synthesis using iScript (Bio-Rad, 170-8891). Samples were analyzed using SYBR Green Supermix (Bio-Rad, 170-8882) reactions run in a CFX96 Optical Reaction Module (Bio-Rad). Expression for each gene was calculated using the formula [62] (Etarget)– ΔCPtarget (control-sample)/(Eref)– ΔCPref (control-sample) and normalized to a reference housekeeping gene (At1g13320). Primer for RT-qPCR analysis were designed by QuantPrime program. Primer sequences are listed in Table S2.

### GUS staining

7-day-old (D8) seedlings were treated with mock (80% ethanol), 50µM IAA and 1µM BL in liquid 0.5xLS on plate and collected after 3h, similar to sample preparation for RT-qPCR. GUS staining was performed as previously described [63] using 1mM Ferri/Ferro concentration for 1.5h at 37°C in the dark. Seedlings were mounted on glass slides in 50% glycerol. Images were taken using Leica microscope and whole seedling images were reconstructed using MosaicJ feature of Fiji plugin from National Institutes of Health ImageJ software (rsb.info.nih.gov).

### ChIP assay

Seven-day-old (D8) seedlings (BES1_PRO_::BES1:GFP, ARF5_PRO_::ARF5:GFP and *bes1-D* x ARF5_PRO_::ARF5:GFP) were treated for 3h with 80% ethanol (mock) or 50µM IAA in liquid 0.5xLS media on plate and crosslinked in 1% formaldehyde under vacuum on ice. Cross-linking was stopped by infiltrating in 0.125M room temperature glycine solution. Seedlings were subsequently frozen in liquid nitrogen and ground to a fine powder with mortar and pestle. Samples were resuspended in nuclei extraction buffer [0.25 M Suc, 100 mM MOPS, pH 7.6, 10 mM MgCl2, 5% Dextran T-40, 2.5% Ficoll, 20 mM b-mercaptoethanol, and mini-Complete Proteinase Inhibitor tablet (Roche Applied Science, 04693124001)], filtered through Miracloth (Calbiochem, 475855), and centrifuged to collect nuclei. Nuclei were lysed with Nuclei lysis buffer (50 mM Tris-HCl, pH 8, 10 mM EDTA, and 1% SDS). ChIP dilution buffer was added (1.1% Triton X-100, 1.2 mM EDTA, 16.7 mM Tris, pH 8.0, 167 mM NaCl, and 0.01% SDS), and chromatin was fragmented using Biorupter sonicator (Fisher Scientific, Bioruptor® UCD-200). An aliquot of fragmented chromatin served as an input control for qPCR analysis, and the remainder was subjected to immunoprecipitation. Dynabeads protein A (Invitrogen, 100-02D) coupled with anti-GFP (Ab290, Abcam) antibody were used to enrich for ARF5_PRO_::ARF5:GFP or BES1_PRO_::BES1:GFP containing chromatin fragments. Samples were washed and eluted off of Dynabeads using nuclei lysis buffer, and cross-links were reversed by incubating with 300 mM NaCl. DNA was purified using a PCR clean-up kit (Qiagen, 28104). Low Adhesive Dnase/RNase free tubes (Bioplastics, B74030) were used for all the procedures. ChIP-qPCR assay data was normalized to housekeeping gene (At1g13320) coding sequence, and results are expressed as ratios of qPCR signal to the antibody IP of wild-type samples (Figure 1) or IP without antibody in reporter lines (Figure S1). ChIP-qPCR results represent the average of at least 2-4 independent biological replicates. Primers for ChIP-qPCR analysis are listed in Supplemental Table S2.

**Figure 1.**
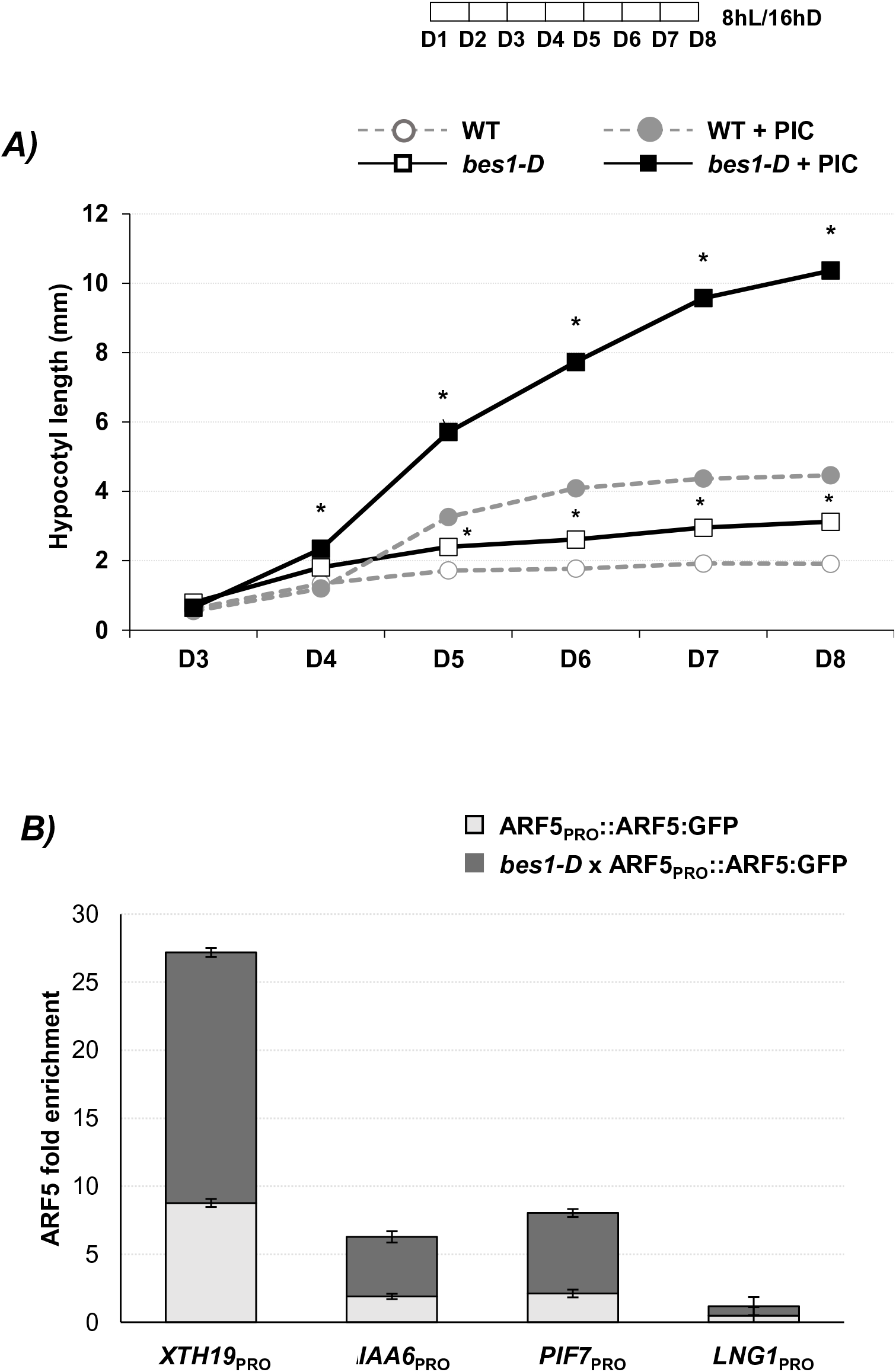
BES1 increases auxin sensitivity and DNA binding by ARF5. **(A)** Hypocotyl response of wild-type (WT) and *bes1-D* gain-of-function mutant seedlings to the synthetic auxin picloram (PIC). Open symbols are seedlings exposed to control treatments, and closed symbols are hormone treated samples. *bes1-D* was significantly taller than WT on D4-under mock and treated conditions. *bes1-D* was significantly longer than WT on D4-8 Asterisks indicate p value< 0.001 **(B)** Binding by ARF5 on several promoters was enhanced in *bes1-D* mutants under mock condition. Bars represent the mean of four biological replicates.

## Results

### BES1 sensitizes hypocotyl to exogenous application of auxin

BRs enhance seedling sensitivity to auxin [22]. To test the extent to which this effect of BR on auxin response is mediated by BES1, we exposed seedlings with wild-type or constitutively active BES1 to the synthetic auxin picloram (Figure1A). As previously described, *bes1-D* hypocotyls were longer than those of wild-type seedlings (Figure 1A; [13]); however, the application of synthetic auxin, picloram, strongly exaggerated the difference in hypocotyl length between *bes1-D* mutant and wild-type plants. Based on previous results using the *SAUR15* promoter [30], we hypothesized that increased BES1 activity led to auxin hypersensitivity by enhancing DNA binding of ARF5 to promoters with bipartite-type *cis*-elements.

Using the large list of BES1 targets generated by ChIP-seq and microarray analysis [16], we focused on the subset of targets with putative bipartite elements in their promoters. Ten genes were selected based on the following criteria: being targets of ARF5 or BES1 [16, 31], having evidence of their expression regulated by auxin or BRs [21, 32], and functional information and/or mutant phenotypes. We validated these candidate genes and the bipartite elements in their promoters with ChIP assays in the previously characterized ARF5_PRO_::ARF5:GFP [31] and BES1_PRO_::BES1:GFP [12] lines. From our initial list, *XYLOGLUCAN ENDOTRANSGLUCOSYLASE/HYDROLASE 19* (*XTH19)*, *INDOLE-3-ACETIC ACID 6 (IAA6)*, *LONGIFOLIA1 (LNG1)* and *PHYTOCHROME-INTERACTING FACTOR7 (PIF7)* were found to be the strongest candidates for further analysis (Figure S1). To test whether BES1 enhances DNA-binding of ARF5, we crossed ARF5_PRO_::ARF5:GFP line to *bes1-D* mutant. Increased BES1 activity did increase DNA binding by ARF5 in most cases (Figure 1B).

### BES1 and BEH4 are major regulators of BR responses in seedling stage

BES1 belongs to a family of six genes [13], with BES1 and BZR1 the best-characterized members [15, 16]. BEH1-4 undergo BR-induced phosphorylation status changes similar to BES1 and BZR1 [13]. The functional redundancy of this transcription family was documented recently in trait robustness [33], however their role in BR pathways remains poorly understood. BEH4, the most recent member of the family, acts redundantly with BES1 to regulate hypocotyl length in skotomorphogenic seedlings [33]. To investigate whether other members of the BES1/BZR1/BEH gene family contribute to auxin sensitivity, we analyzed T-DNA insertion alleles for each member of the BES1/BZR1/BEH family [33]. In these T-DNA insertion lines, we measured hypocotyl elongation in the absence or presence of BL or picloram. Hypocotyl response to BL was modestly reduced only in *bes1-2*, *bzr1-2* and *beh4-1* single mutants (Figure S2), while the response to picloram was not affected significantly in any of single mutants (Figure S3). Based on these findings, we selected *bes1-2 beh4-1* and *bes1-2 bzr1-2* double mutants for further investigation. *bes1-2 beh4-1* double mutants are dwarfs that resemble known mutants with compromised BR synthesis or signaling (Figure 2A). They also show dramatically reduced response to BL and significantly reduced response to picloram (Figures 2B and C). In addition, rosettes of *bes1-2 beh4-1* resembled *bin2-D* mutants where activity of the entire BES1/BZR1/BEH family should be suppressed (Figure 2A). In contrast, *bes1-2 bzr1-2* double mutants had an essentially wild-type response to picloram, but significant reduction in BR sensitivity (Figure S4).

**Figure 2.**
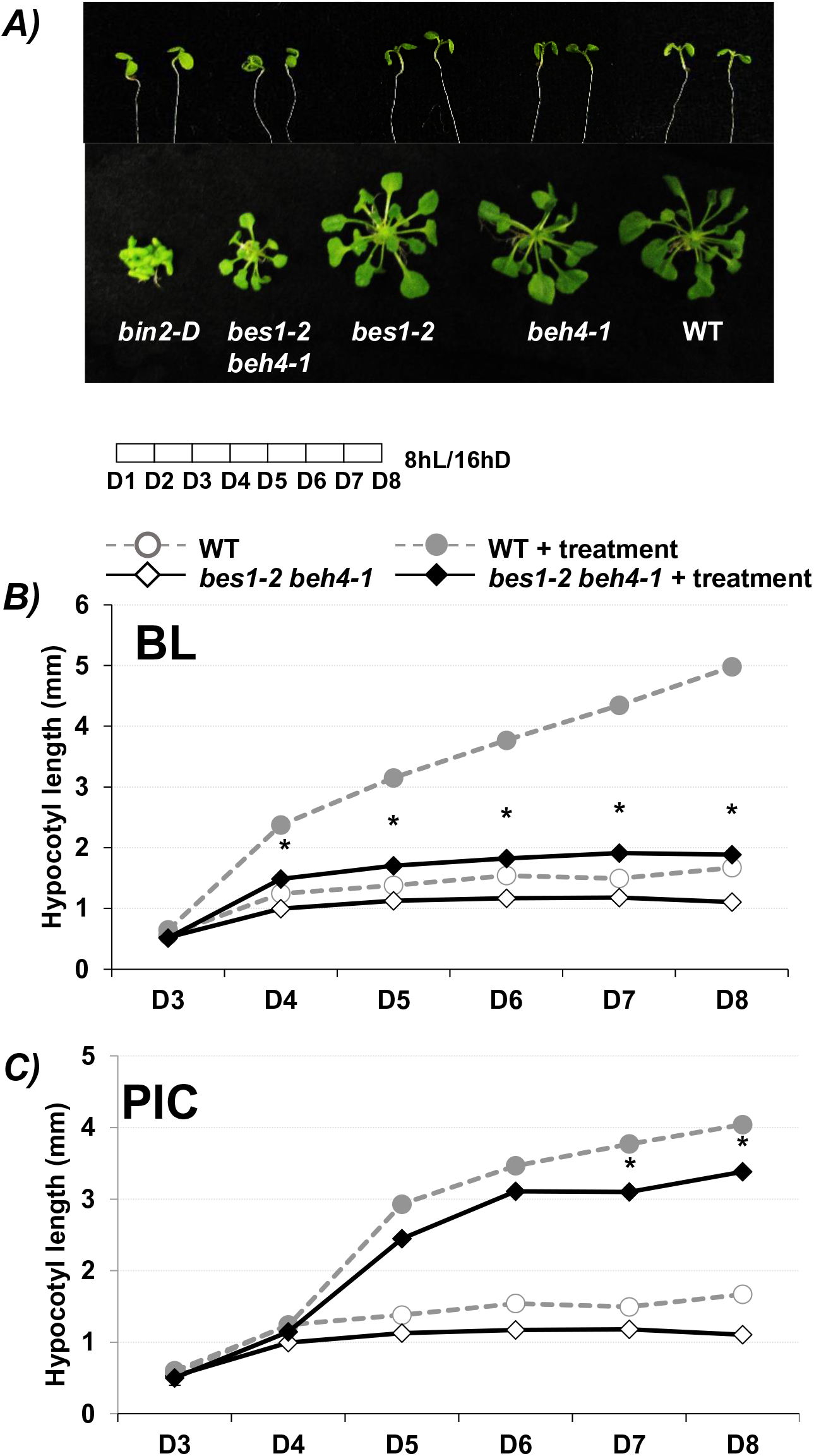
BEH4 acts redundantly with BES1 to regulate auxin and BR seedling responses. **(A)** 7-day-old seedling (upper panel) and 6-weeks rosette (lower panel) phenotype of wild-type, *bin2-D, bes1-2 beh4*, *bes1-2* and *beh4-1* mutants grown in short day conditions. Hypocotyl response of wild-type (WT) and *bes1-2 beh4-1* mutant seedlings to **(B)** brassinolide (BL) and **(C)** picloram (PIC), grown as indicated in Figure1. Open symbols are seedlings exposed to control treatments, and closed symbols are hormone treated samples. *bes1-2 beh4-1* was significantly shorter than WT on D4-8 (p value< 0.01) under control and brassinolide, and only on D7-8 (p value< 0.05) picloram conditions. WT and *bes1 beh4-1* control samples are the same and repeated for reference in panel B and C.

### BES1 status affects bipartite gene expression

We used RT-qPCR to investigate the effect of BES1 on bipartite target gene expression (*XTH19*, *IAA6*, *LNG1* and *PIF7*). Single mutants of *bes1-2* and *beh4-1* did not have significant effect on target gene expression (Figure S5), consistent with their weak physiological phenotypes (Figures S2 and S3). However, in *bes1-2 beh4-1* double mutant the expression of *XTH19* and *IAA6* was attenuated, and the expression of *LNG1* was modestly decreased. In contrast, the expression of *PIF7* was induced in the double mutant. As expected, in *bes1-D* the opposite tendencies were observed for all genes (Figures 3A). Both BR and auxin dramatically induced the expression of *XTH19* and *IAA6*; while both hormones modestly repressed the expression of *PIF7* (Figure 3B). The hormone responsiveness of those genes was affected by BES1/BEH4 status (Figure 3C).

**Figure 3.**
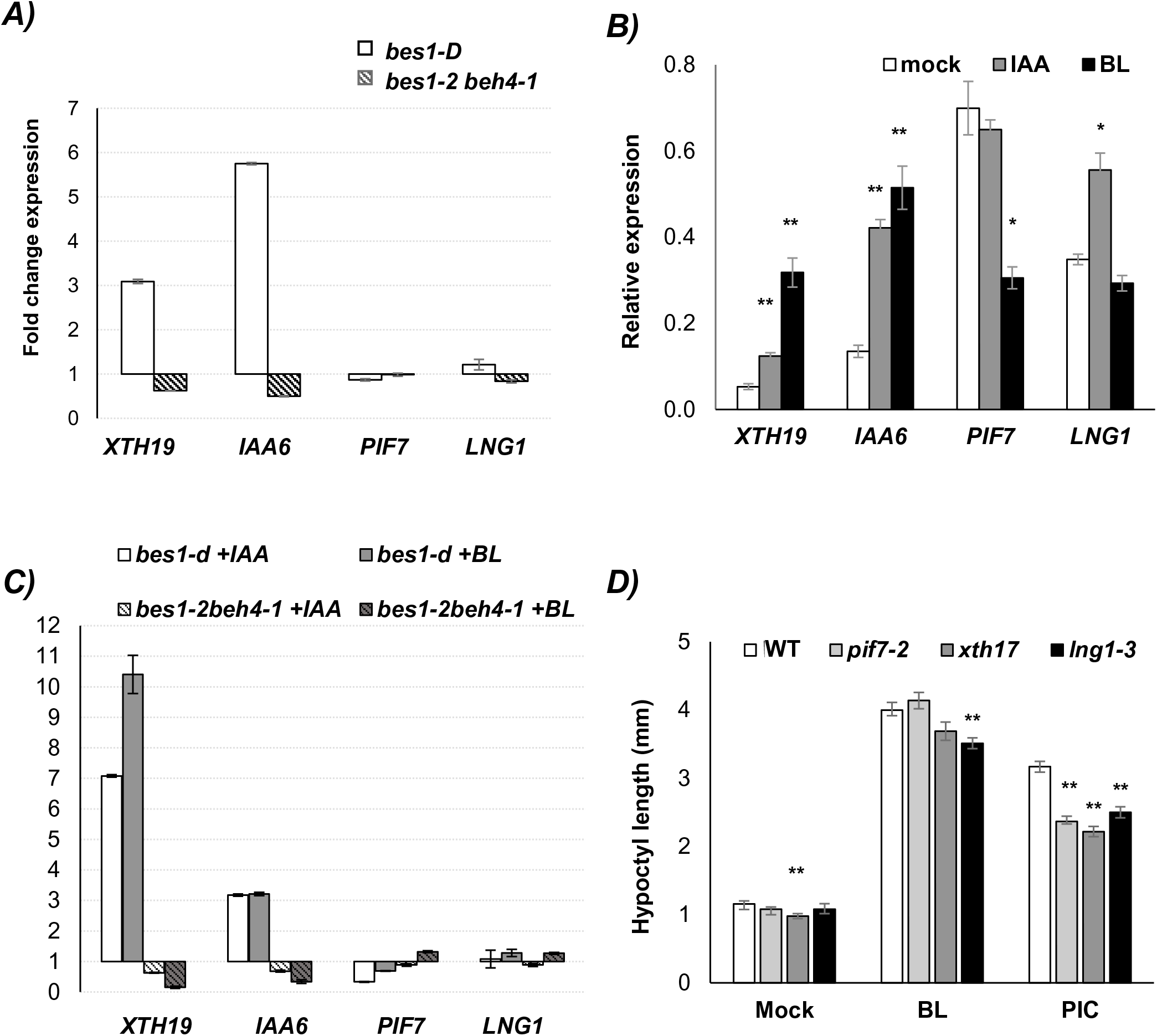
Expression of ARF5-BES1 targets is hormone sensitive and important for growth. **(A)** BES1 and BEH4 status affects the expression of ARF5 targets. RT-qPCR analysis of bipartite target gene expression in 7-day-old seedlings without any treatment. The fold change expression of *XTH19*, *IAA6*, *PIF7* and *LNG1* is shown in *bes1-D* and *bes1-2 beh4-1* double mutants relative to WT. **(B)** Hormone responsiveness of target genes in WT under control (mock), 3h of IAA or BL treatment. **(C)** Hormone responsiveness of target genes is altered in *bes1-D* and *bes1-2 beh4-1* double mutant. Means of three biological replicates (relative expression value normalized to housekeeping gene) is shown ± SE in A-C. **(D)** ARF5 target genes are important for growth. Hypocotyl response of 7-day-old WT, *pif7-2*, *xth17* and *lng1-3* mutant seedlings under control (mock), brassinolide (BL) or picloram (PIC) treatments. *\**:p value< 0.05; *\*\**:p value< 0.01 in comparisons between mock and treated samples in (B) and WT vs. mutant phenotypes in (D).

We further investigated the role of bipartite target genes in BR and auxin induced elongation responses. We analyzed single knockout alleles of *xth17*, *lng1-3* and *pif7-2*. Since *XTH19* had no available loss-of-function mutant, we analyzed a mutant in *XTH17*, part of the same *XTH17-20* gene cluster whose members act redundantly with one another [34, 35]. *XTH17* promoter also carried a conserved bipartite element, and its expression followed a similar pattern as *XTH19* in our RT-qPCR analysis (Figure 4B). The hypocotyls of all three mutants tended to be shorter than those in wild-type seedlings in control (mock) conditions. In the presence of picloram, these effects became more obvious (Figure 3D).

**Figure 4.**
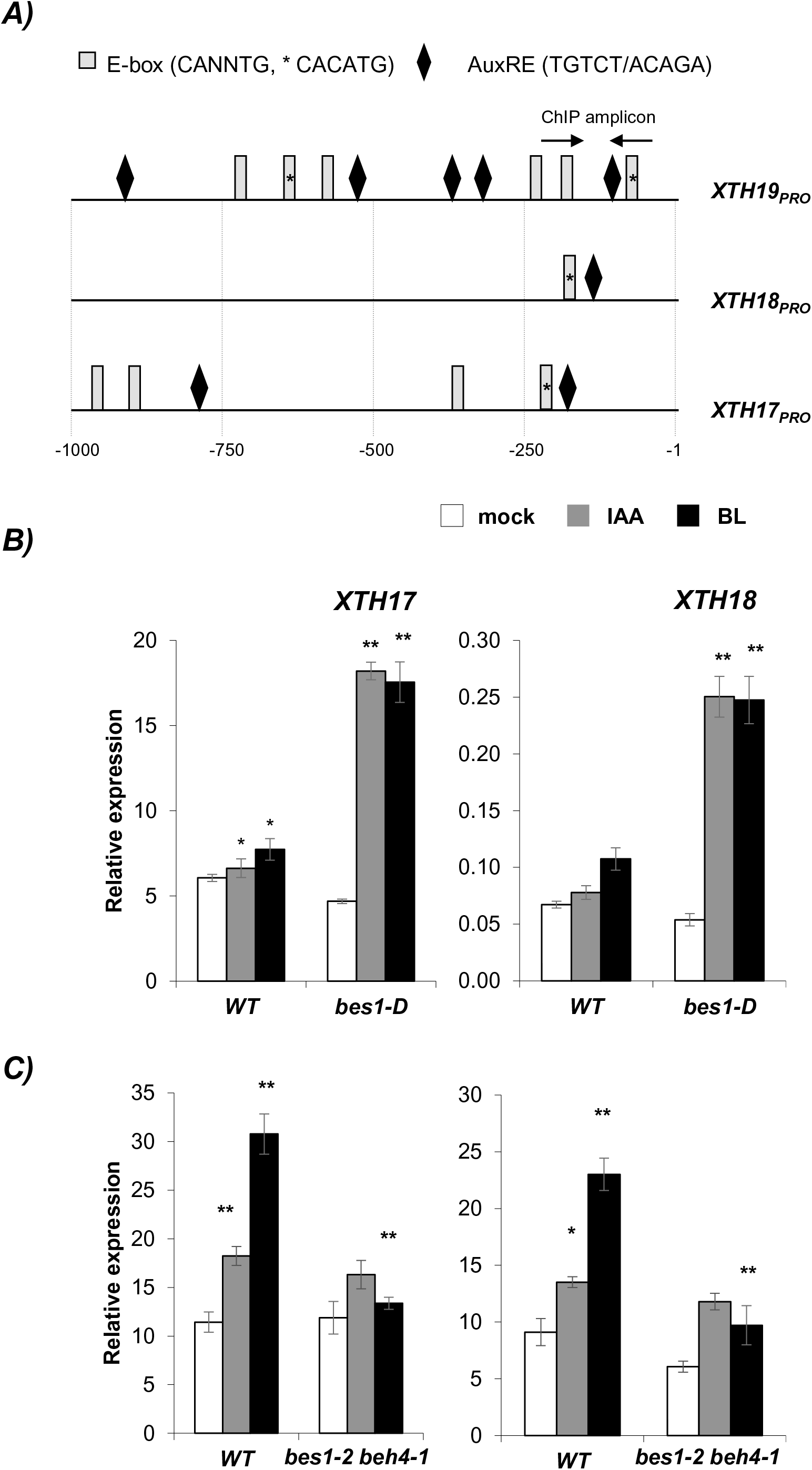
Other members of *XTH19* gene clade with bipartite promoter structure exhibit similar hormone responsiveness in a BES1-dependent manner. **(A)** Promoter architecture of *XTH* genes. One kilobase upstream of the transcriptional start site is shown. qPCR analysis of several *XTH* gene expression in 7-day-old seedlings in response to control (mock), IAA or BL in **(A)** WT, **(B)** *bes1-D* and **(C)** *bes1-2 beh4-1* double mutant. Means of three biological replicates (relative expression value normalized to housekeeping gene) is shown ± SE. *\*\** refers to p value< 0.01, *\** to p value< 0.05.

### Evolutionary conservation of bipartite elements predicts hormone responsiveness of promoters

*XTH19*, *IAA6* and *LNG1*, as well as the previously characterized bipartite target gene *SAUR15*, belong to large gene families. In the case of *XTH19*, we observed that the expression of at least three genes in the same clade (*XTH17-19*) were similarly affected by BES1, as well as treatment with BR or auxin (Figures 4B and C). We reasoned that these similar transcriptional patterns across paralogs might be due to shared promoter architecture. Previously, it was documented that *XTH17-20* cluster has a specific motif conservation within the promoter region [34]. We found that this conserved motif includes the bipartite element we found in *XTH19* promoter (Figure 4A). Similar promoter architecture is also observed for group of so-called *SAUR* class 2 and 3 that are expressed in hypocotyls [36]. 17 out of 30 genes in this group harbor bipartite elements within 250bp upstream of the transcription start, and all of these have been shown to be regulated by BR and auxin (Table S1). We predicted that the absence of bipartite element would compromise hormone responsiveness of target genes. In fact, no hormonal responses to either BL or IAA was observed for several *XTHs* closely related to *XTH17-19* cluster, such as *XTH14*, *31* and *32*, in which either HUD or AuxRE elements were lost. *XTH31* expression was still responsive to BL, likely due to a retained HUD element at −900bp. *XTH26* has conserved bipartite element and was included as a positive control. Despite its low expression levels, *XTH26* was repressed by both hormones.

In previous work focused on characterization of the *XTH17-20* cluster, Vissenber and colleagues generated a set of transgenic lines of promoter truncations fused to GUS. We tested full-length (XTH19_PRO-1.1kb_::GUS) and 300bp (XTH19_PRO-0.3kb_::GUS) truncated promoter reporter lines in our conditions. Seedlings of full-length XTH19_PRO-1.1kb_::GUS line exhibited weak staining in the hypocotyl area and root tip in control (mock) conditions, 3h of IAA and BL treatment intensified the blue staining specifically in hypocotyl and petiole tissues. Interestingly the XTH19_PRO-0.3kb_::GUS truncation, in which a portion of the bipartite elements is deleted, completely abolished reporter gene expression and hormone responsiveness (Figure S6). These data suggest that bipartite elements in *XTH19* promoter are important for its proper expression level, pattern and hormone-sensitivity.

## Discussion

During photomorphogenesis, seedling growth is shaped by a complex interacting networks of plant hormones to insure the seedling architecture and growth are synchronized to environmental conditions. In this study, we provide molecular mechanism for how two well-characterized regulators of photomorphogenesis, auxin and BRs, converge in a tunable bipartite transcriptional module to promote growth.

We have shown that BES1 and BEH4, but not BZR1, play a major role in BR and auxin responsiveness in young seedlings. These results are consistent with a recent study that found that BEH4 acts redundantly with BES1 in controlling hypocotyl length robustness in dark-grown seedlings [33]. BES1 is also temperature sensitive, a trait regulated by auxin [37, 38]. It is likely that other family members, such as BZR1, play growth-promoting roles at other stages or under other conditions. Consistent with this hypothesis, ChIP-seq analyses of BES1 and BZR1, performed in two-weeks old seedlings and leaves of adult plants, respectively, found a large overlap in target promoters [15, 16]. BES1 and BZR1 show differential interaction with the chaperone HSP90 [37] which may facilitate interactions with distinct cell type- or stage-specific partners.

Cooperative binding and activity of BR-regulated transcription factors appear to be a recurrent motif in plant signaling. Our group previously found that the interaction between auxin and BR relies on two cis-elements, a bipartite element that contains AuxRE and HUD-type E-box elements bound by ARF5 and BES1, respectively [30]. Here, we show that specific mutations of E-boxes/AuxRE elements during evolution or disruption of bipartite target XTH19_PRO_ had direct effect on gene expression level and patterns (Figure 5 and S6). A similar mechanism of cooperative binding has been proposed for BZR1 and ARF6 [29]. BES1 also interacts with the bZIP transcription factors HAT1 and HAT3 to co-repress the BR biosynthetic gene *DWF4* [39]. In addition, both BES1 and BZR1 interact with the bHLH transcription factor PIF4 and bind to E-box elements to co-activate target promoters [40].

**Figure 5.**
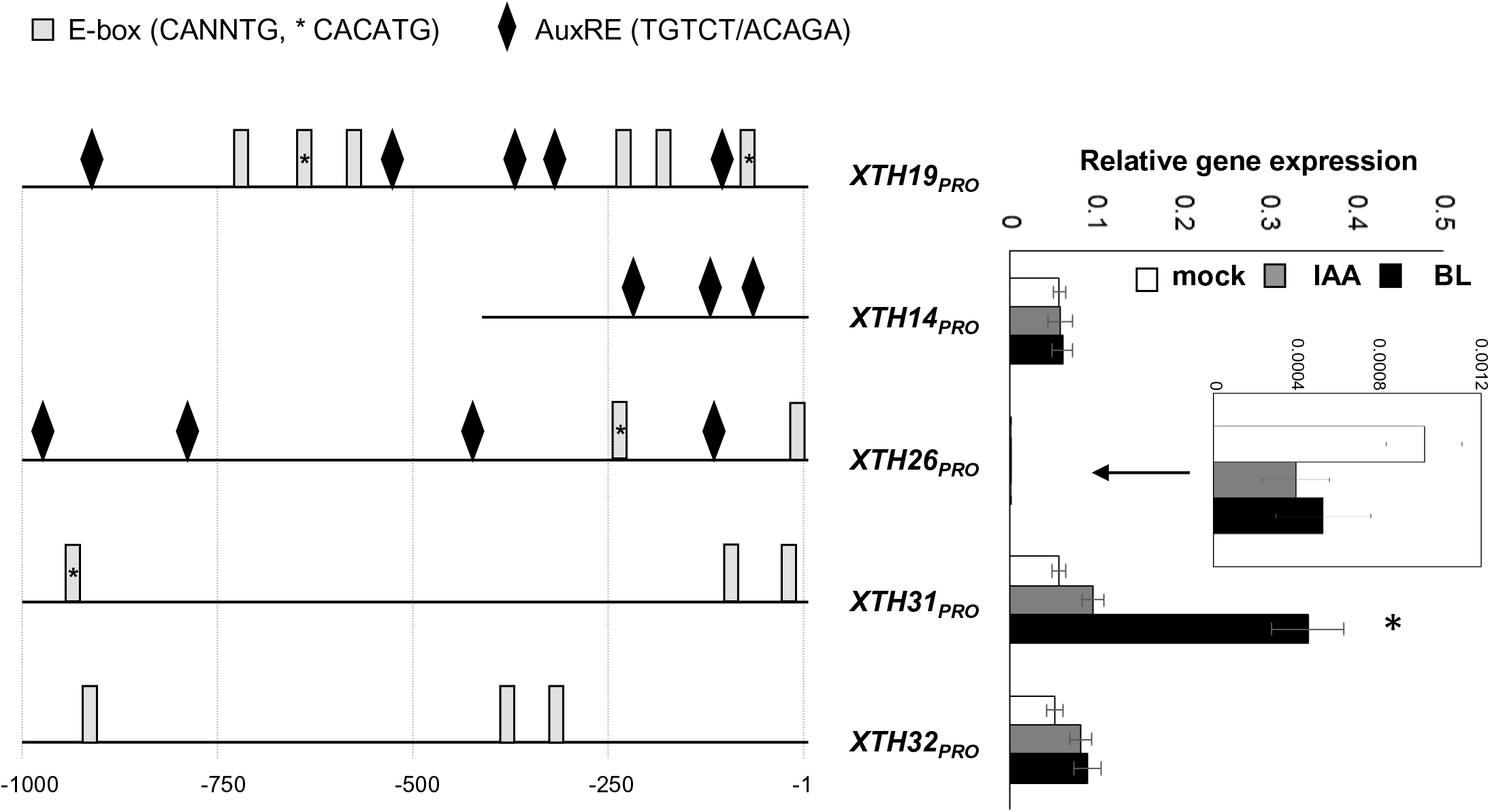
Promoter architecture predicts hormone sensitivity. **Left panel:** Promoter architecture of non-bipartite *XTH* genes. One kilobase upstream of the transcriptional start site is shown. **Right panel:** RT-qPCR analysis of selected non-bipartite *XTH* gene expression in 7-day-old wild-type in response to 3h of control (mock), 50µM IAA or 1µM BL (relative expression value normalized to housekeeping gene in WT). *: p value< 0.05 of significant difference between treated and mock samples.

Target genes with bipartite promoters identified in this study are known regulators of plant growth. *IAA6*, also called *SHORT HYPOCOTYL1*, encodes a co-repressor of auxin signaling that is also involved in negative feedback [41]. PIF7 is as major regulator of shade responses that directly induces auxin biosynthesis genes such as members of the *YUCCA* family [42]. Dominant *iaa6/shy1-1D* and loss-of-function *pif7* mutants exhibit short hypocotyl phenotypes [43, 44] (Figure 3D). *LNG1* encodes a protein localized to cortical microtubules (cMT) [45] that was initially identified as a dominant mutant with exaggerated elongation of petioles via unidirectional cell elongation [46, 47].

The switch between different cMT orientations can be triggered by various endogenous and exogenous signals that are known to modulate growth, including light and various hormones [48–50]. Actin and cMT proper orientation and localization are required for both auxin and BR-mediated cell elongation [51–53]. In addition, Sasidharan and colleagues demonstrated that genetic and pharmacological disruption of cMTs affects a shade-specific subset of *XTH* genes (*XTH17* and *XTH19*) expression as a result of auxin re-distribution [52]. XTHs, similar to expansins, are cell wall modifiers that induce cell expansion. Overexpression of *XTH18, XTH19* and *XTH20* (all part of the bipartite *XTH* clade) stimulated hypocotyl growth in early developmental stage of Arabidopsis seedlings [35], while similarly loss-of-function of *XTH17* results in inhibition of hypocotyl growth (Figure 3D). In addition, the functionally redundant *LNG3* and *LNG4* genes regulate turgor-driven polar cell elongation through activation of *XTH17* and *XTH24* [54]. BES1/BZR1 and PIFs were also implicated in hypocotyl elongation during thermomorphogenesis via regulation of *LNG1* and *LNG2* [55–57]. We found that *LNG1* and *XTH19* are bipartite targets of the ARF5-BES1/BEH4 module rapidly induced by both auxin and BRs (Figures 3 and 4) further connecting reorientation of cMTs and cell wall loosening.

We propose that the ARF5-BES1/BEH4 transcriptional hub rapidly and coordinately modulates a suite of growth control genes (Figure 6). This module can serve as an integration point for external signals such as shade and temperature to tune internal growth program and adjust it to a changing environment. For example, exposure to shade could rewire this growth network by increasing the expression/activity of bHLH transcription factors that interact with BES1, and, in this way, change the composition and thereby the targets of the auxin/BR transcriptional complex. Future studies are needed to fully understand how each potential transcriptional complex impacts target gene selectivity and growth dynamics in a tissue- and developmental stage-specific manner.

**Figure 6.**
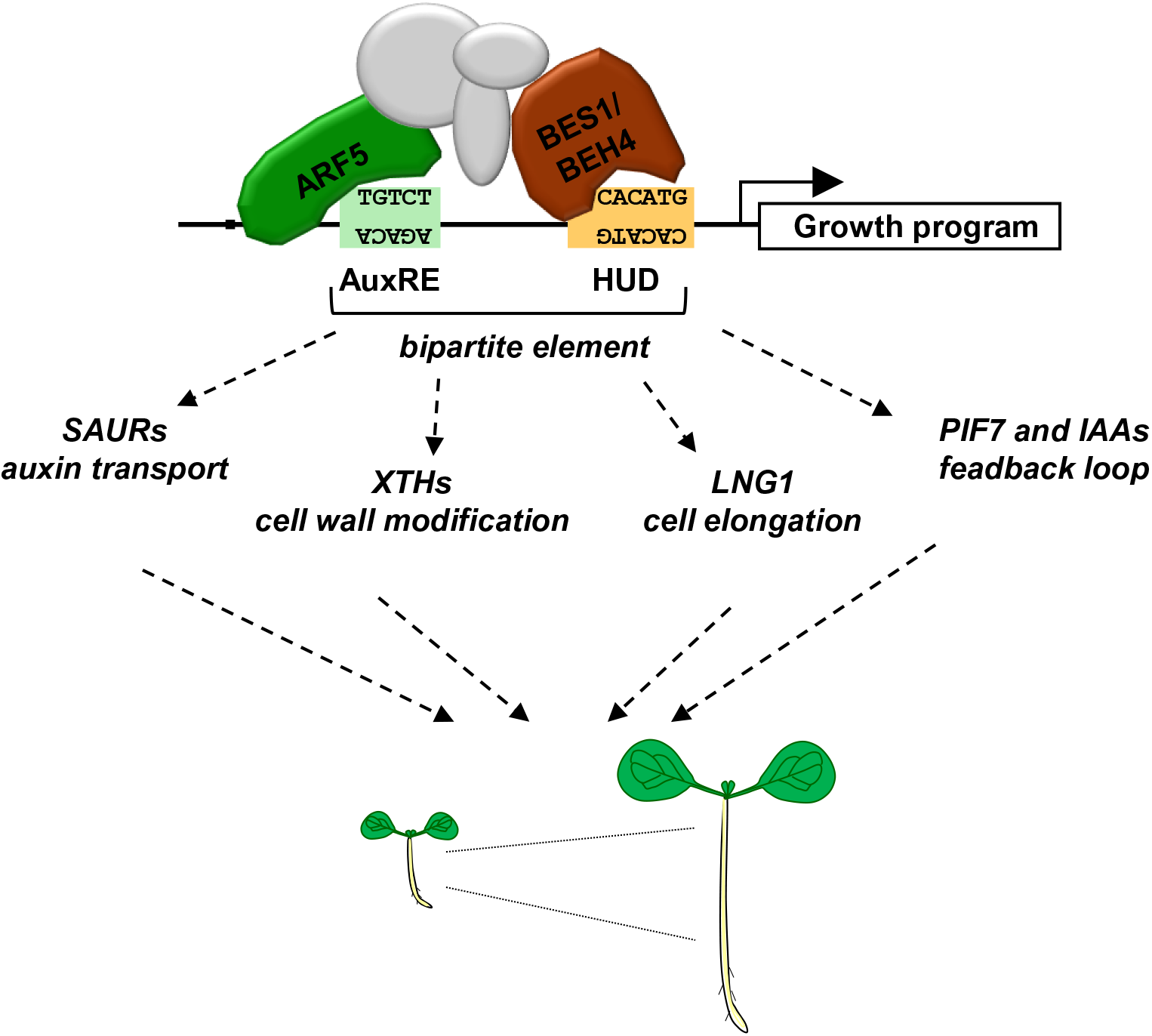
ARF5-BES1-BEH4 transcriptional hub acts as a molecular switch to integrate signals. Bipartite elements allow ARF5, BES1 and BEH4 to work together as a transcriptional module to connect expression of a suite of growth-promoting genes to specific environmental conditions.

## Acknowledgements

This work was supported by the National Institutes of Health (R01-GM107084). We thank Dr. Jennifer Lachowiec and Dr. Christine Quietsch for providing seed stocks before publication, as well as Dr. Dolf Weijers, Dr. Zhi-Yong Wang, Dr. Peter Quail, Dr. Kazuhiko Nishitani, Dr. Ronald Pierik and Dr. Jaime F Martinez-Garcia for sharing published resources.

## Authors Contribution

AG and JLN conceived and designed the experiments. AG collected and analyzed the data, as well as preparing the figures. AG and JLN wrote the manuscript.

## Supporting Information

**Figure S1:** BES1 and ARF5 share target genes

**Figure S2:** BES1-BZR1-BEH family regulates seedling establishment during photomorphogenesis in the presence of brassinosteroids.

**Figure S3:** BES1-BZR1-BEH family single loss-of-function mutants do not affect seedling establishment during photomorphogenesis in the presence of picloram.

**Figure S4:** BZR1 contribution to auxin response in seedling stage.

**Figure S5:** Single loss-of-function of BES1 and BEH4 do not alter bipartite gene expression.

**Figure S6:** Bipartite elements are required for proper expression pattern and hormonal activation of XTH19_PRO_

**Table S1.** List of SAUR genes with bipartite elements.

**Table S2.** List and nucleotide sequence of primers used in this work.

