## Supplemental Data for "Auxin promotion of seedling growth via ARF5 is dependent on the brassinosteroid-regulated transcription factors BES1 and BEH4"

### Supplementary Materials and Methods

#### Plant Crosses and Phenotypes Analysis

*bes1-2*, *beh4-1* and *bzr1-2* single mutants are T-DNA insertion lines and were PCR genotyped using gene specific and T-DNA specific primers. *bes1-D* single point mutation was genotyped using dCAP method: PCR amplification of the region containing the single point mutation and BglII digestion where only mutant fragment is restriction sensitive. *bes1-D* x $ARF5_{PRO}$::ARF5:GFP line was generated by crossing *bes1-D* single mutant and $ARF5_{PRO}$::ARF5:GFP reporter line. $F_2$ generation was screened for $BASTA^R$ (for $ARF5_{PRO}$::ARF5:GFP) seedlings and genotyped with specific BglII-sensitive dCAP primers (for *bes1-D*). Genotyping primers for *bes1-D*, *bes1-2*, *beh4* and *bzr1-2* are indicated in Table S1.

### References

### Supplementary figure legends

**Figure S1. BES1 and ARF5 share target genes.**

ChIP-qPCR assay of **(A)** BES1 and **(B)** ARF5 enrichment on target promoters in $BES1_{PRO}$::BES1:GFP and ARF5 $_{PRO}$::ARF5:GFP reporter lines respectively. Bars are mean of two biological replicates, normalized to housekeeping gene and represented relative to no antibody (α-GFP) negative control samples of reporter line. ACT2-CDS is shown as a non-specific reference. Samples were harvested on D8 according to the scheme.

**Figure S2. BES1-BZR1-BEH family regulates seedling establishment during photomorphogenesis in the presence of brassinosteroids.**
Hypocotyl response of wild-type (WT) and *bes1-2*, *bzr1-2*, *beh1-1*, *beh2-1*, *beh3-1* and *beh4-1* mutant seedlings to brassinolide, grown as indicated in upper pannel. Open symbols indicate control and closed symbols, treated samples. * refers to *t*-test p value< 0.01.

**Figure S3. BES1-BZR1-BEH family regulates seedling establishment during photomorphogenesis in the presence of auxin.**
Hypocotyl response of wild-type (WT) and *bes1-2, bzr1-2, beh1-1, beh2-1, beh3-1* and *beh4-1* single loss-of-function mutant seedlings to picloram (PIC), grown as indicated in upper pannel. Open symbols indicate control and closed symbols, treated samples. Only *bzr1-D* mutant response to picloram is statistically significant (p value ≤0.01) compared to WT. * refers to *t*-test p value< 0.001.

**Figure S4. BZR1 contribution to auxin response in seedling stage. (A)** and **(B)** Hypocotyl response of wild-type (WT) and *bes1-2 bzr1-2* mutant to brassinolide (BL) and picloram (PIC). Open symbols indicate control and closed symbols, treated samples. WT and *bes1 bzr1-2* control samples are the same and repeated for reference in panel B and C. *bes1-2 bzr1-2* was significantly shorter than WT on D5-8 (* p value< 0.01) under brassinolide conditions only.

**Figure S5. Single loss-of-function of BES1 and BEH4 do not alter bipartite gene expression.**
RT-qPCR analysis of bipartite target gene expression in 7-day-old seedlings in response to control (mock), IAA or BL in wild-type, **(A)** *bes1-2* and **(B)** *beh4-1* mutant. Means of three biological replicates (relative expression value normalized to housekeeping gene) is shown ± SE. ** refers to p value< 0.01; * p value< 0.05 of significant differences between mock and treated samples in WT, and corresponding sample in WT and mutant as indicated in Figure (A).

**Figure S6. Bipartite elements are required for proper expression pattern and hormonal activation of XTH19$_{PRO}$ in hypocotyl. (A)** Schematic representation of *XTH19*$_{PRO}$ and its modifications. **(B)** GUS staining of 7-day-old transgenic seedlings from reporter lines XTH19$_{PRO}$

$_{-1.1kb}$::GUS and XTH19$_{PRO-0.3kb}$::GUS after 3h of mock, IAA and BL. The blue color marks the expression domains of promoter reporters.

**Table S1. List of SAUR genes with bipartite elements. Classification of the genes based on Sun et al 2016.**

| | Gene ID | Gene discription | Bipartite elements |
|---|---|---|---|
| class 1 | AT4G38840 | SAUR-like auxin-responsive protein family 14 | + |
| | AT1G75580 | SAUR-like auxin-responsive protein family 51 | + |
| class 2 | AT2G21210 | SAUR-like auxin-responsive protein family 6 | + |
| | AT4G38850 | SAUR-like auxin-responsive protein family 15 | + |
| | AT5G18030 | SAUR-like auxin-responsive protein family 21 | + |
| | AT4G34760 | SAUR-like auxin-responsive protein family 50 | + |
| | AT1G29430 | SAUR-like auxin-responsive protein family 62 | + |
| | AT1G29450 | SAUR-like auxin-responsive protein family 64 | + |
| | AT1G29510 | SAUR-like auxin-responsive protein family 67 | + |
| class 3 | AT2G21200 | SAUR-like auxin-responsive protein family 7 | + |
| | AT2G18010 | SAUR-like auxin-responsive protein family 10 | + |
| | AT4G38825 | SAUR-like auxin-responsive protein family 13 | + |
| | AT5G18050 | SAUR-like auxin-responsive protein family 22 | + |
| | AT4G13790 | SAUR-like auxin-responsive protein family 25 | + |
| | AT3G03850 | SAUR-like auxin-responsive protein family 26 | + |
| | AT1G29440 | SAUR-like auxin-responsive protein family 63 | + |
| | AT1G29460 | SAUR-like auxin-responsive protein family 65 | + |
| | AT1G29500 | SAUR-like auxin-responsive protein family 66 | + |
| | AT1G29490 | SAUR-like auxin-responsive protein family 68 | + |
| class4 | AT4G34790 | SAUR-like auxin-responsive protein family 3 | + |
| | AT2G16580 | SAUR-like auxin-responsive protein family 8 | + |
| | AT2G28085 | SAUR-like auxin-responsive protein family 42 | + |
| | AT1G29420 | SAUR-like auxin-responsive protein family 61 | + |
| | AT3G03847 | SAUR-like auxin-responsive protein family 73 | + |
| others | AT5G66260 | SAUR-like auxin-responsive protein family 11 | + |

### Table S2. List and nucleotide sequence of primers used in this work.

| Purpose | Gene | ID | Forward | Reverse |
|---|---|---|---|---|
| ChIP-qPCR | *XTH19*$_{PRO}$ | AT4G30290 | TTGGATTCACTTGGCAACAG | TGGTGCATGTGCTTTTGTCT |
| | *IAA6*$_{PRO}$ | AT1G52830 | TCTCTCATGTGACCGACCAA | AGTTGCTCCAACAAGATCCAA |
| | *PIF7*$_{PRO}$ | AT5G61270 | ACACAAACAATCCTTTGCCA | CTCAAAAATGGTCCCTGCAA |
| | *LNG1*$_{PRO}$ | AT5G15580 | TTGGATTGAAAGCAAAATAAAGG | TGATGAAAGTGATGCACATGAA |
| RT-qPCR | *XTH17* | AT1G65310 | CCGAGTCAATTGGAGTTCCGTTCC | TTCCCAGAGACTCGCGTAAAGC |
| | *XTH18* | AT4G30280 | CCGAGTCAATTGGAGTTCCGTTCC | TTCCCAGAGACTCGCGTAAAGC |
| | *XTH19* | AT4G30290 | ATGAATGCCGAGTCACGTGGAG | TCGCATATAGCCTCATTGGTTGC |
| | *XTH14* | AT4G25820 | TCACCGCCTACTACCTATCGTC | AGGATGTCCTGTGCGATTTCCC |
| | *XTH26* | AT4G28850 | CCCGACCAATGGTTTCCACAAC | GAGTTCCATCGACGAACCACAC |
| | *XTH31* | AT3G44990 | TGGGAGTGGGTTCAAGTCTCTTCG | TGAAGCCTGGTTGGAGCTTAATGG |
| | *XTH32* | AT2G36870 | CTGGCTTGATAGAACCTCAGGAAG | CCAAAGTAGCCTGATCTGAATGGC |
| | *IAA6* | AT1G52830 | TTCGGCTGTCTTGGCATAGGAG | TCCGACGAGCATCCAGTCTCTATC |
| | *PIF7* | AT5G61270 | ATTCACAACGAGTCCGAAAGGAG | GCAGCTTCTGAAGTGTTCTCATCC |
| | *LNG1* | AT5G15580 | AGGAGTCCAGACTTGGGAATGAGG | TGGGCTGCTCCACTGTAACTTTC |
| | *LNG2* | AT3G02170 | AAGAGGAGGATGCCTTACGTTCTG | ACGGATCTGCAGAAGCTGGTAG |
| | *BES1* | AT1G19350 | GGCTGGTTTAACTCAAATCAACGG | TCCGTCAGACGTCATCTTCTTCG |
| | *BEH1* | At3g50750 | TTTGTCTTGAAGCTGGTTGGATCG | TTCTGTTGGTCGAGAACCCTTTC |
| | *BEH2* | At4g36780 | TATCCAACAGTGCGCCTGTGAC | AGTTTCCGCTTCGAACCACGAG |
| | *BEH3* | At4g18890 | TGCAATGAAGCTGGTTGGACTG | TCCATTGGTTTGCATCCCTTGC |
| | *BEH4* | AT1G78700 | GCACTCTGTAACGAAGCTG | TGGCTGATAGGAAGAGCA |
| | *BZR1* | AT1G75080 | CGGTGAACCAAATAACAACATGTC | GATAGATTCCCAGTTAGGCAAC |
| | *BZR1 FL* | AT1G75080 | TGGCCGTCGCGAACCATGACTTCGGATGGAGCTAC | TCCCATTCGCGATCAACCACGAGCCTTCCC |
| | *ACT2* | AT3G18780 | CAAGCTGTTCTCTCCTTGTACGC | ACCAGAATCCAGCACAATACCG |
| | *UNC21* | AT5G25760 | GACCAAGATATTCCATCCTA | GTTAAGAGGACTGTCCG |
| | *HK* | AT1G13320 | AACGTGGCCAAAATGATGC | AACCGCTTGGTCGACTATCG |
| Genotyping | *bes1-2* | AT1G19350 | WT-CAGTCAGGACAAAAGTAAGCACTC | CCCTAAAGGTCACTTTCTCCG |
| | | | T-DNA- AACGTCCGCAATGTGTTATTA | |
| | *beh1-1* | At3g50750 | WT - CATTGGACTCGATTCTCGAAG | GGGAGTTTTCTCGGTGAGATC |
| | | | SAIL LB- TAGCATCTGAATTTCATAACCA | |
| | *beh2-1* | At4g36780 | WT-CATTGGACTCGATTCTCGAAG | GGGAGTTTTCTCGGTGAGATC |
| | | | SAIL LB- TAGCATCTGAATTTCATAACCA | |
| | *beh3-1* | At4g18890 | WT-ACCTCGATCGTTGTACAAAGG | AGCCTGAGCACGTGTTAACAC |
| | | | LB3- TAGCATCTGAATTTCATAACCAATCTCGATACAC | |
| | *beh4-1* | AT1G78700 | WT-ACCACAATAGCCAAATTGCTG | TGAAATCCAAATCGCAAGATC |

| | | | | |
|---|---|---|---|---|
| | | | LB3-TAGCATCTGAATTTCATAACCAATCTCGATACAC | |
| | *bes1-D* | AT1G19350 | CCTACTCATCATCGCCAGTTCCAAGATC | TTGAAAGCTTATCCAATGACCA |
| | *bzr1-2* | AT1G75080 | CATCAGTCTACGTTCACACAATC | WT-AACCAATCAAACATCAATCAATCA |
| | | | | T-DNA-ATATTGACCATCATACTCATTGC |

WT refers to wild-type; HK to housekeeping genes used to qPCR normalizations; BZR1-FL to full-length CDS.

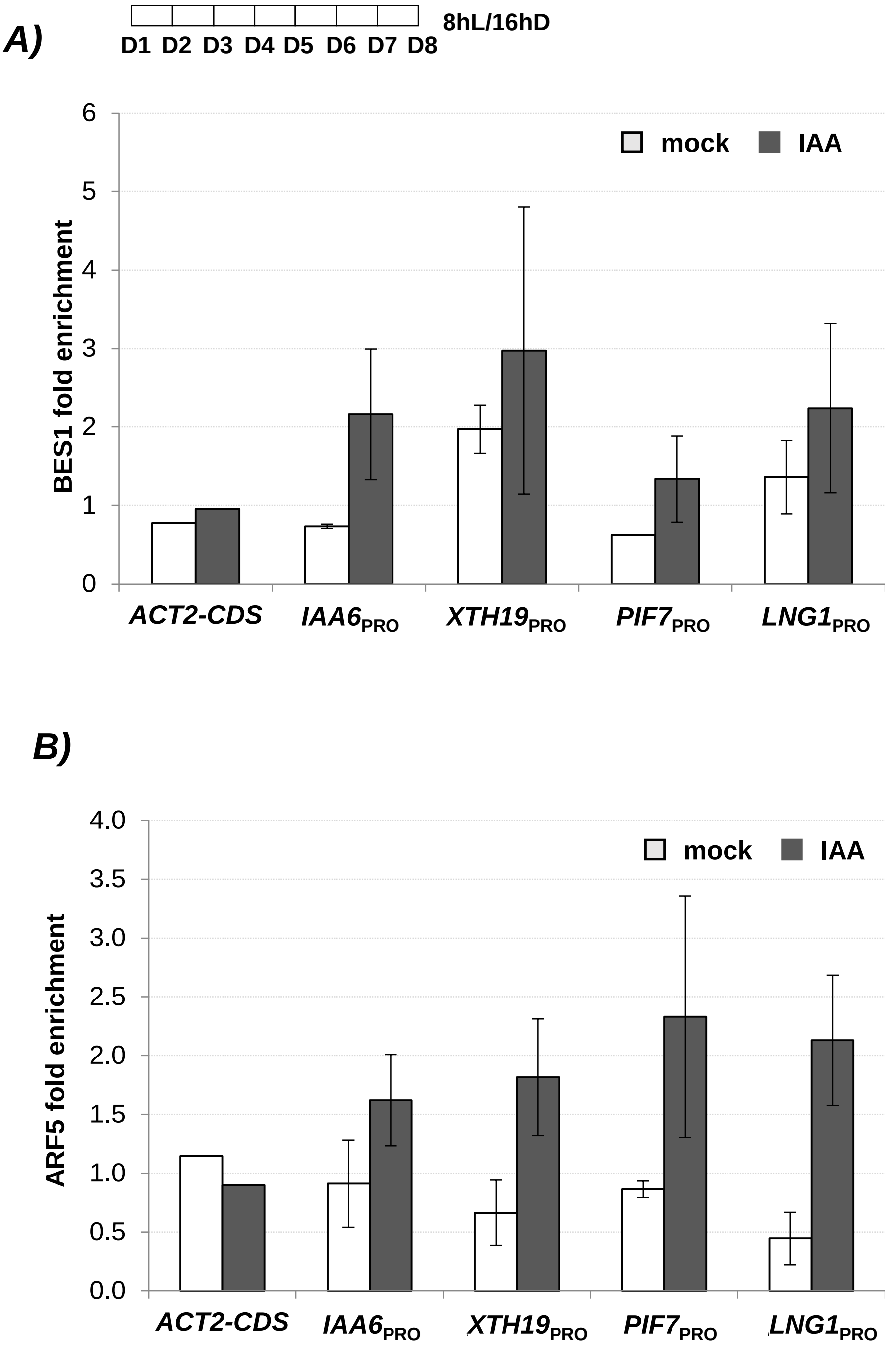


**Figure S1. BES1 and ARF5 share target genes.**

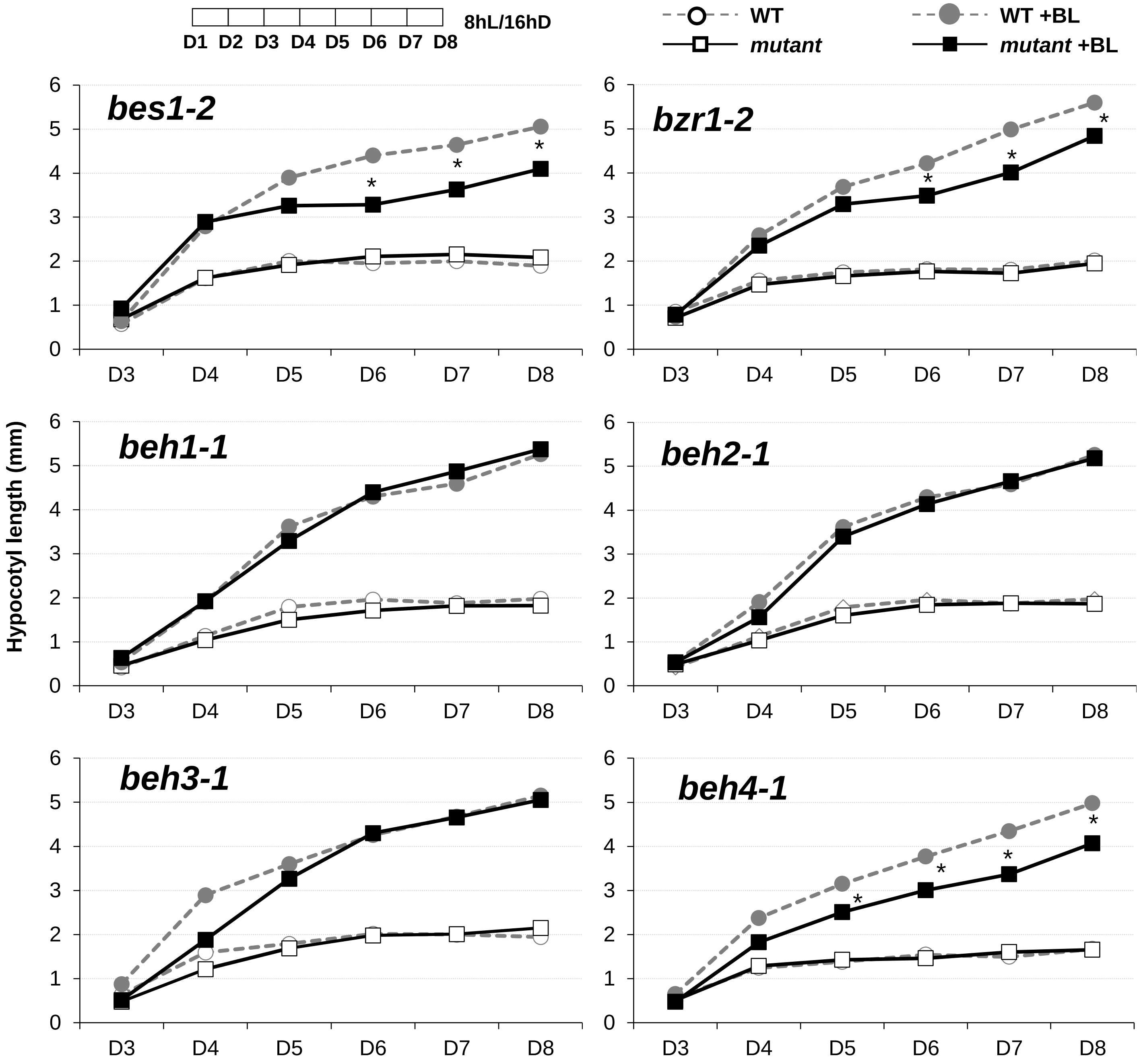


**Figure S2. BES1-BZR1-BEH family regulates seedling establishment during photomorphogenesis in the presence of brassinosteroids.**

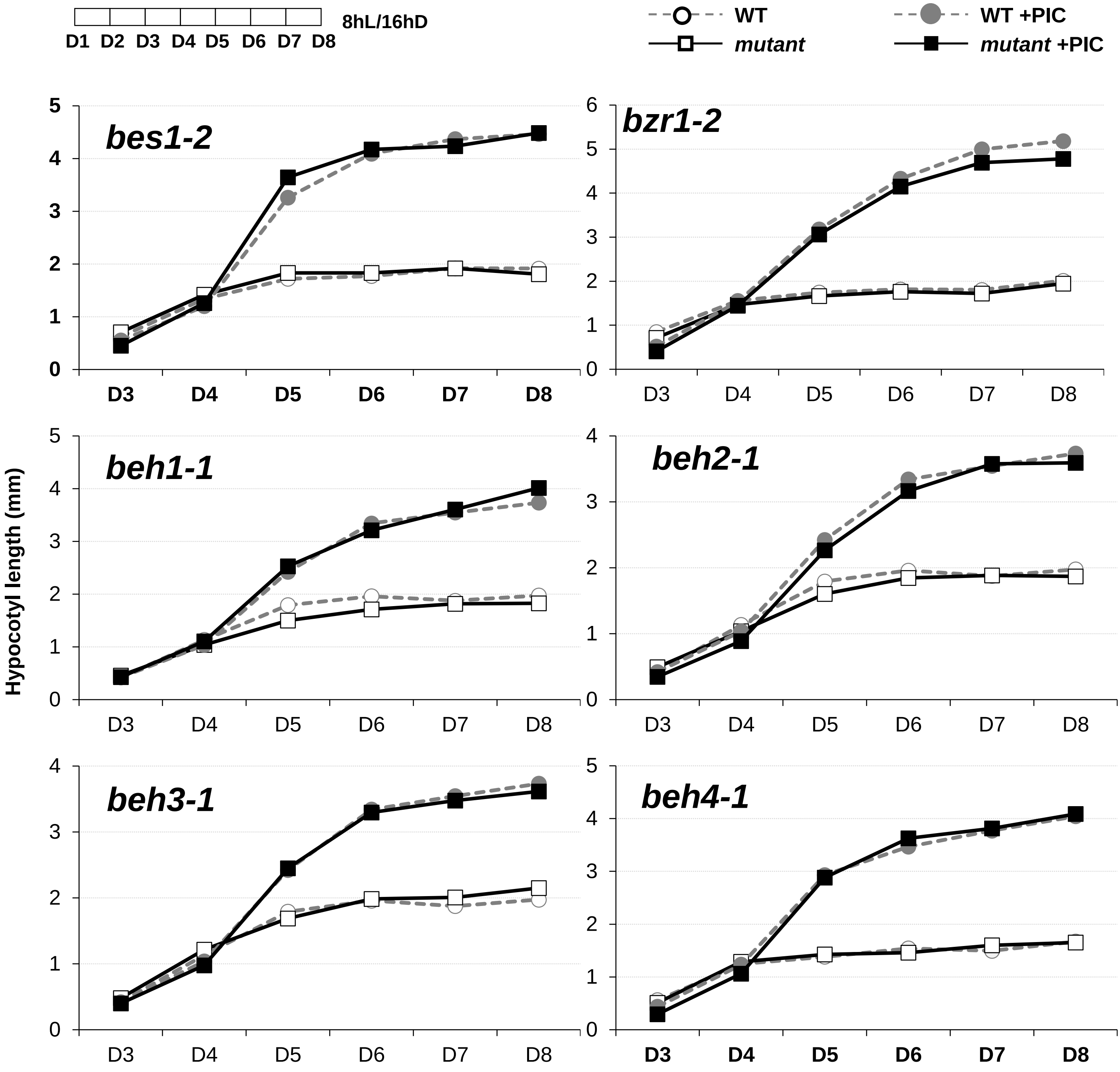


**Figure S3. BES1-BZR1-BEH family single loss-of-function mutants do not affect seedling establishment during photomorphogenesis in the presence of picloram.**

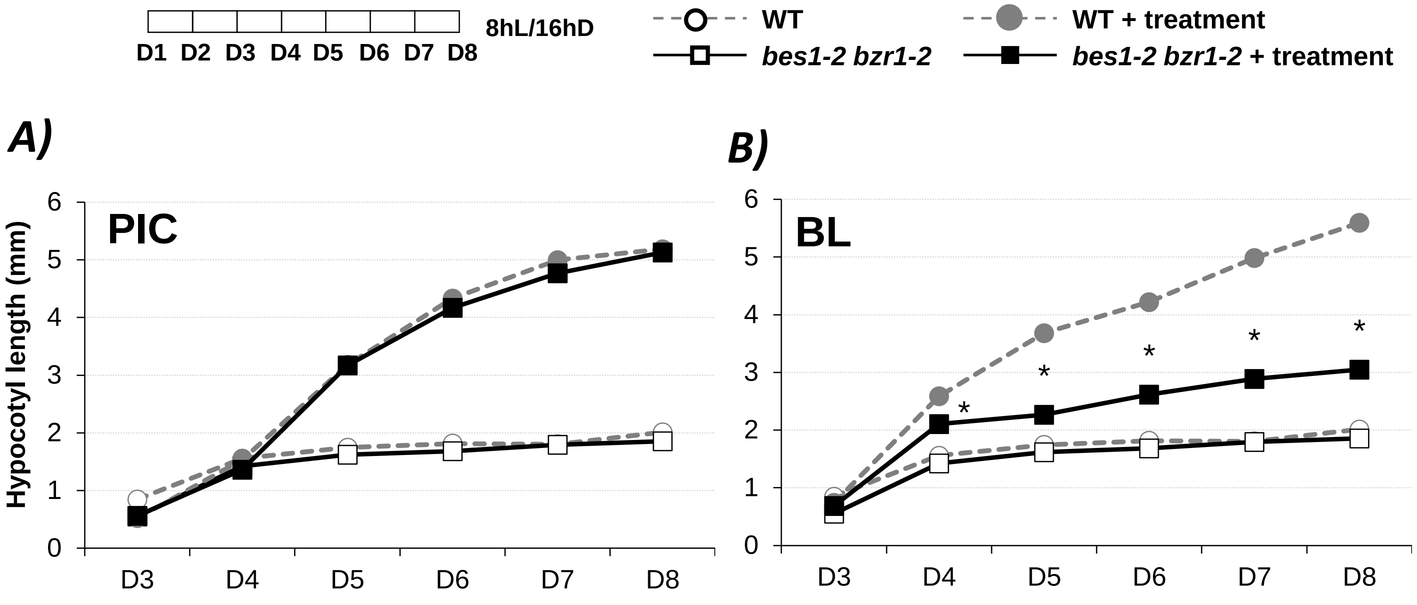


**Figure S4. BZR1 contribution to auxin response in seedling stage.**

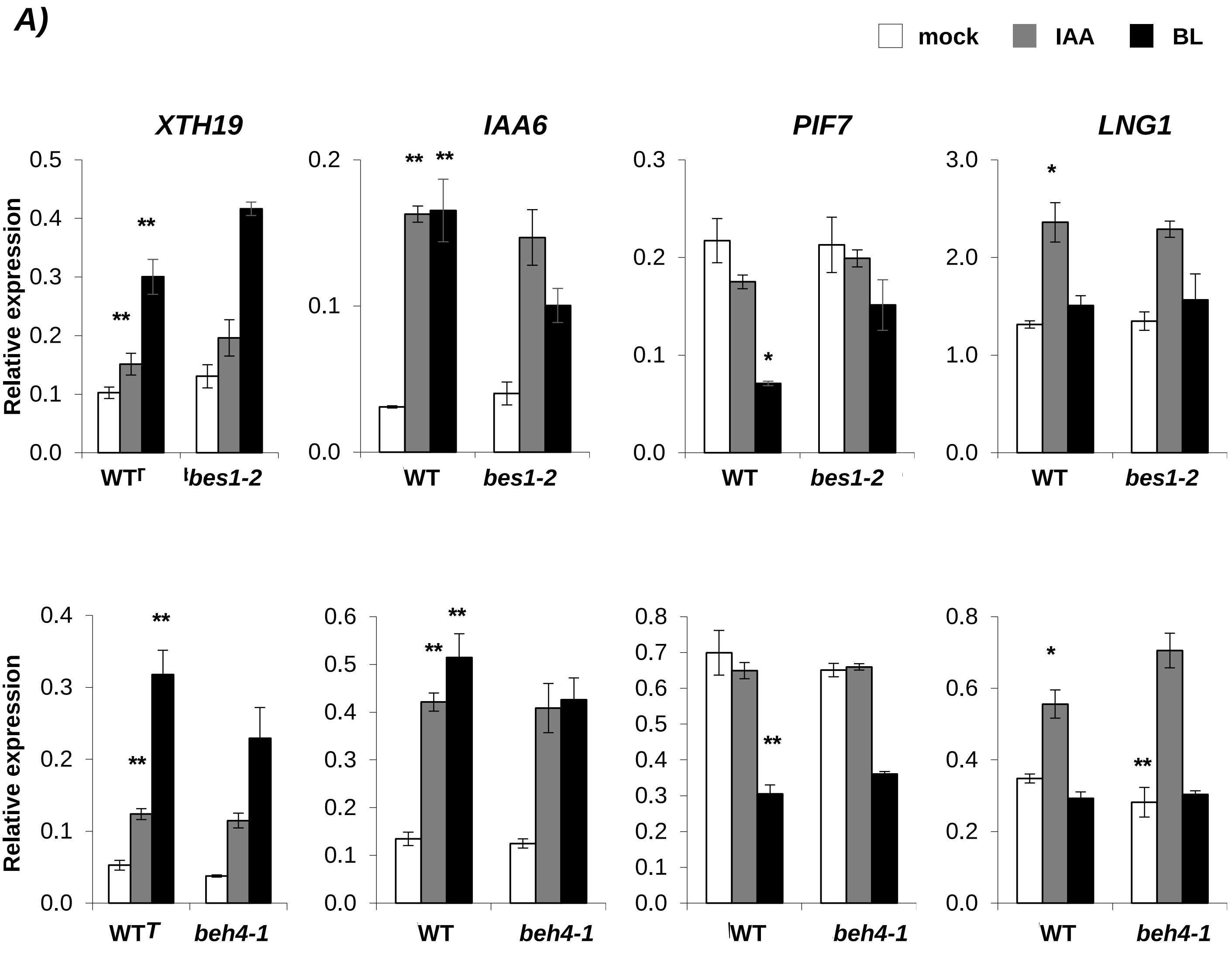


**Figure S5. Single loss-of-function of BES1 and BEH4 do not alter bipartite gene expression.**

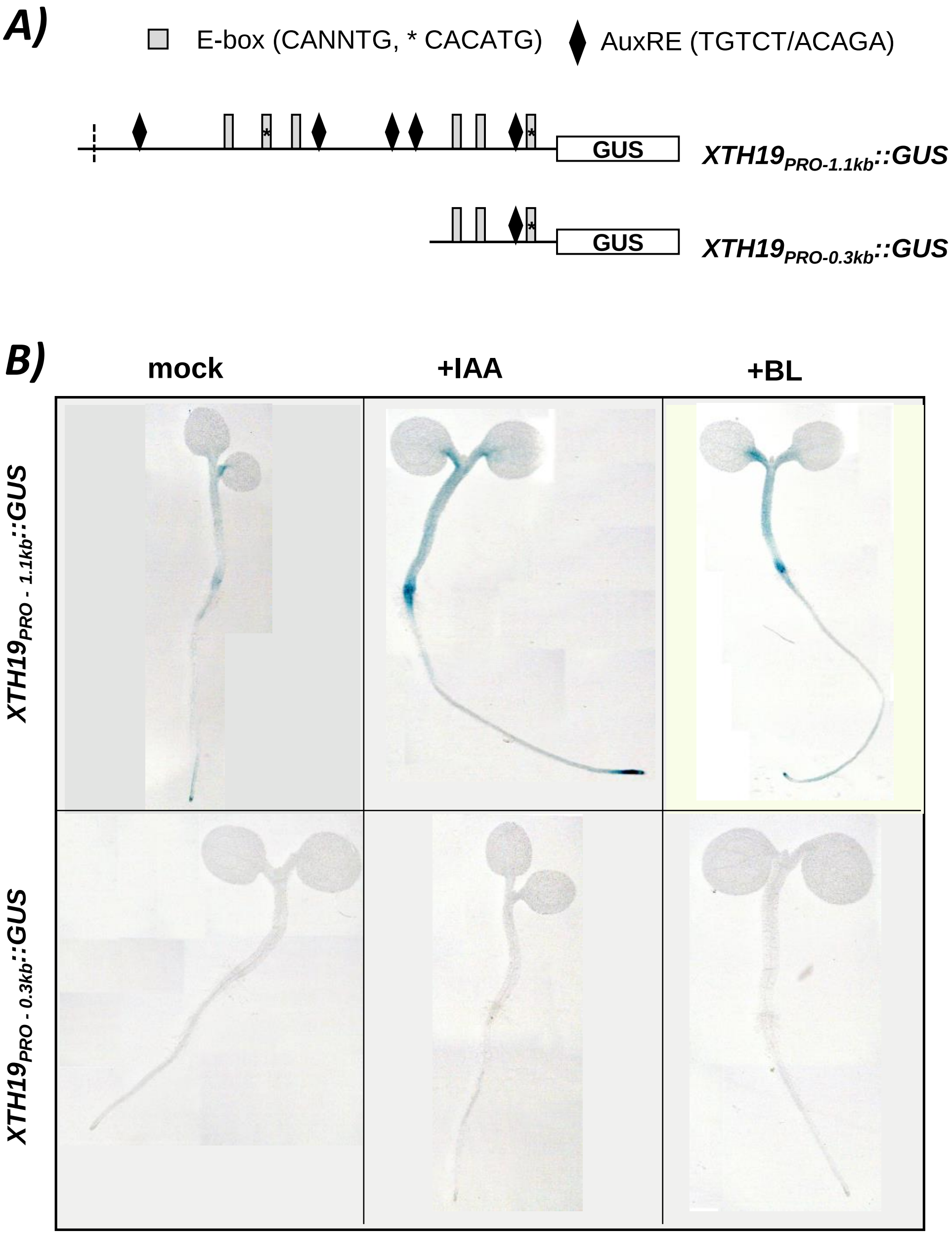


**Figure S6. Biparrtite elements are required for proper expression pattern and hormonal activation of *XTH19*$_{PRO}$ in hypocotyl.**
